# High-Throughput Human Gut Immune Co-Culture Model for Evaluating Inflammatory Bowel Disease Anti-Inflammatory Therapies

**DOI:** 10.1101/2025.05.14.654072

**Authors:** Swetha Peddibhotla, Lauren A. Boone, Earnest Taylor, Bryan E McQueen, Elizabeth M. Boazak

## Abstract

Current treatments for inflammatory bowel disease (IBD) are often ineffective long-term, as many patients ultimately become unresponsive to anti-inflammatory drugs. The need for improved therapeutics is urgent. Animal models utilized for drug development are limited by interspecies variability and poor translatability. However, most in vitro models lack the sophistication to model the key interplay of the immune system with the intestinal epithelium in line with the known role of the immune system in the etiology of the disease.

To address this gap, we developed a primary intestinal epithelial cell co-culture system to incorporate elements of innate immune signaling. This system models immune-epithelial interactions using RepliGut^®^ - Planar Transverse Colon cultured on a Transwell™ system with THP-1 derived macrophages in a receiver compartment of a 96-well plate. Epithelial barrier integrity and cell viability were maintained in co-culture with unstimulated macrophages. However, similar to the pathology associated with IBD, epithelial integrity was compromised in co-culture with LPS + IFN-γ pre-stimulated macrophages as evidenced by declining TEER and cell viability and increased inflammatory cytokine release. Cotreatment with anti-inflammatory IBD therapeutics adalimumab or tofacitinib mitigated these effects, demonstrating the model’s ability to replicate key inflammatory responses and prevention.

Reproducibility and scalability of the model system further position the model for high-throughput screening of anti-inflammatory drugs, improving drug discovery, and accelerating the translation of new IBD therapies into clinical practice.

**Highlights:**

- Co-culture model: RepliGut^®^ - Planar Transverse Colon with THP-1 derived macrophages
- High throughput and human-relevant model
- “Healthy” co-culture resembling healthy intestine
- “Inflamed” co-culture mimicking IBD innate inflammatory signaling
- Potential to screen anti-inflammatory drugs relevant to IBD

**Graphical Abstract:** Gut-immune co-culture model simulating healthy and inflamed intestine. The immune co-culture model consists of mature differentiated primary human transverse colon epithelial cells cultured on a 96-well Transwell^®^ plate with macrophage differentiated THP-1 cells (THP-1m) cultured in the receiver plate. In this configuration, the THP-1m are located basally to the epithelial cells, allowing for apical treatment in the transwell and basal treatment in the receiver plate. In the unstimulated state, intestinal cells and immune cells maintain a stable co-culture. Upon stimulation with LPS and IFN-y, both cell types initiate an inflammatory response that results in release of cytokines, loss of intestinal barrier integrity, and cytotoxicity.

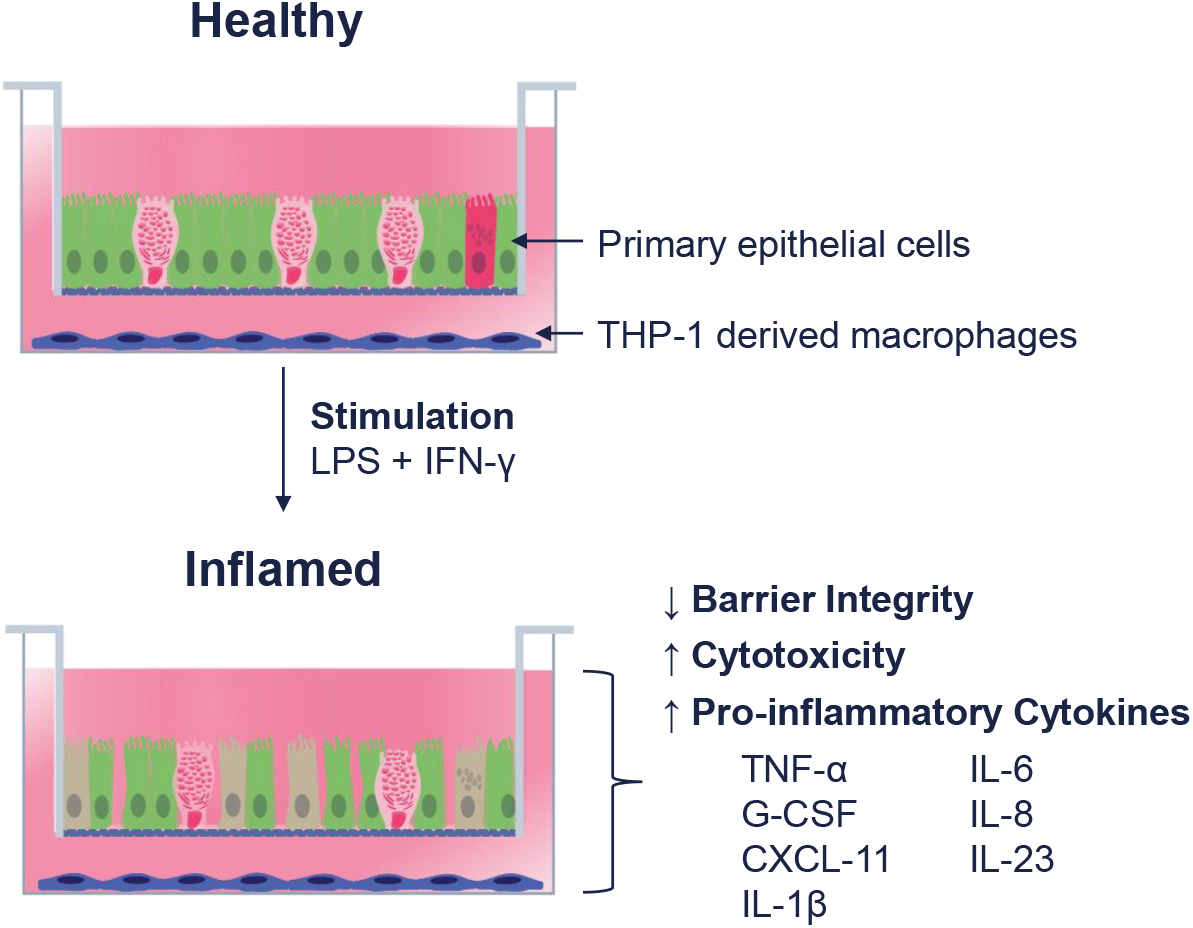

## 1. Introduction

Inflammatory bowel disease (IBD) encompasses two chronic conditions—Crohn’s disease (CD) and ulcerative colitis (UC)—that result from chronic inflammation in the gastrointestinal (GI) tract (Szigethy et al., 2010). Crohn’s disease can affect any part of the GI tract, while UC primarily affects the colon. Inflammatory bowel disease prevalence is increasing globally, particularly in North America, Europe, and Australia, with an estimated 3 million cases in the U.S alone (Lin et al., 2024). Symptoms associated with IBD include, but are not limited to, abdominal pain, diarrhea, and fatigue (Guan, 2019). While current therapies, such as anti-TNF biologics and small molecule JAK inhibitors, have significantly improved patient outcomes, they are not universally effective. A substantial portion of patients exhibit primary non-response, develop secondary loss of response over time, or experience dose-limiting side effects (Sandborn et al., 2022). Furthermore, existing treatments primarily manage inflammation rather than offering a cure, highlighting a critical unmet clinical need. This therapeutic gap is, in part, attributed to the limitations of preclinical models used in drug discovery, which often fail to accurately predict human efficacy and toxicity. Therefore, developing models that better recapitulate the complex human disease state is crucial for identifying novel therapeutic targets and advancing more effective treatments.

The complex pathophysiology of IBD presents significant challenges in developing physiologically relevant preclinical models. Currently, researchers rely on animal models, cell-line derived culture and co-culture models, or a combination (Joshi et al., 2022). However, commonly used models possess significant limitations in recapitulating human IBD. For instance, the widely used dextran sulfate sodium (DSS) mouse model induces colitis primarily through direct epithelial toxicity, largely bypassing the complex, immune-mediated mechanisms central to human IBD pathogenesis, such as T-cell activation and cytokine-driven epithelial dysfunction (Chassaing et al., 2014; Eichele & Kharbanda, 2017). Furthermore, animal models (e.g., mice) exhibit known differences in gut anatomy (Treuting et al., 2012), immune responses (Mestas & Hughes, 2004), and in intestinal epithelial receptor expression, particularly within the TLR family (Pott & Hornef, 2012).

In vitro models offer advantages such as increased throughput and extension to human cells to provide relevant mechanistic insights. However, there are limitations. The Caco-2 cell line, derived from human colorectal adenocarcinoma, is frequently employed as an in vitro model of the human intestine, but lacks the cellular diversity (e.g., secretory cells including goblet cells) and functional differentiation characteristic of the native human gut epithelium (Lennernäs, 2007; Sambuy et al., 2005; Sun et al., 2008, 2008; VanDussen et al., 2015). Caco-2 cells also exhibit altered barrier properties, lack native inflammatory signaling pathways, and have demonstrated altered immune responses compared to primary intestinal epithelial cells (Janssen et al., 2024). These factors make them poorly suited to model the dynamic, cytokine-mediated epithelial injury seen in IBD. Simpler in vitro models using epithelial cells alone, often stimulated with select cytokines, fail to capture the synergistic signaling pathways of the native environment and miss the crucial epithelial-immune cell crosstalk. While co-culture models using Caco-2 cells have been developed (Kämpfer et al., 2017), models incorporating primary intestinal cells have been more challenging to establish due to limited availability of large cell banks from individual donors and inherent run-to-run variability frequently associated with primary cells. These collective limitations underscore the need for more physiologically relevant in vitro systems, particularly those incorporating primary human cells and immune components, that can accurately model the human intestinal epithelial responses to immune-mediated inflammation (Ambrosini et al., 2020; Le et al., 2023; Macedo et al., 2023). Such models are critical for improving our understanding of the innate cellular mechanisms driving IBD and for identifying potential new therapeutic targets.

Therefore, we established a human-relevant high-throughput co-culture model that mimics the innate macrophage-epithelial inflammatory environment of the IBD-affected gut. In this paper, we present the development and characterization of a co-culture model comprised of primary human intestinal epithelial cells (RepliGut^®^ - Planar Transverse Colon) on a 96-well Transwell™ plate with Human Acute Monocytic Leukemia Cell Line 1 (THP-1) derived macrophages (THP-1m) in the receiver compartment. The THP-1 cells were selected for the immune component due to their reproducibility and their ability to differentiate into macrophages which produce key pro-inflammatory cytokines associated with IBD (Sharma et al., 2024). To induce an “inflamed” state, we employed the well-established Toll-Like Receptor 4 (TLR-4) agonist Lipopolysaccharide (LPS) and Interferon-Gamma (IFN-γ), which are known to produce a robust macrophage response (Fultz et al., 1993; Huang et al., 2019; Young & Denovan-Wright, 2024). Co-stimulation with LPS and IFN-γ in macrophages triggers the release of pro-inflammatory cytokines associated with IBD, such as Tumor Necrosis Factor-Alpha (TNF-α), Interleukin-1 Beta (IL-1β), and Interleukin-6 (IL-6) (Kämpfer et al., 2017). To establish the translational relevance of our co-culture model, we evaluated the effects of two clinically established IBD therapeutics: adalimumab, a TNF-α blocking antibody, and tofacitinib, a small molecule JAK inhibitor. By assessing the ability of these drugs to modulate the induced inflammatory response within our system, we aimed to establish a benchmark for the model’s predictive capacity in evaluating anti-inflammatory efficacy. This co-culture model allows researchers to further investigate the complex interactions between primary human epithelial cells and macrophages and provides a high throughput platform to evaluate potential IBD therapies.

## 2. Materials and Methods

### 2.1. Cell Culture

#### 2.1.1. THP-1 Monocyte Expansion

The human leukemic cell line THP-1 was obtained from ATCC (Cat#TIB-202). THP-1 cells were cultured in RPMI-1640 medium (ATCC, Cat#30-2001) supplemented with 8% FBS (R×D systems, Cat#S11150H), 1% Glutamax (Gibco, Cat#35050061), 0.2% Primocin (Invivogen, Cat#ant-pm-2), and 0.055 mM 2-mercaptoethanol (Fisher Scientific, Cat#AC125472500) at 37°C, 5% CO2 for up to 11 passages, in accordance with ATCC provided protocols.

#### 2.1.2. Plating and Differentiation of THP-1 Cells

THP-1 derived macrophages (THP-1m) were generated as follows: THP-1 cells were thawed, counted, and resuspended in THP-1 medium described in (2.1.1), without mercaptoethanol and containing 100 nM Phorbol 12-myristate 13-acetate (PMA, Millipore Sigma, Cat#5.00582). 2.1x10^4^cells per well were seeded in all studies following the density optimization reported in Fig. 2A. After 24 hours, the medium was replaced with THP-1 medium without mercaptoethanol and without PMA, and then cultured for an additional 24 hours. In both steps, medium volume was 100 µL per well. These steps correspond to the timeline steps shown in pink in Fig. 1.

**Figure 1.**
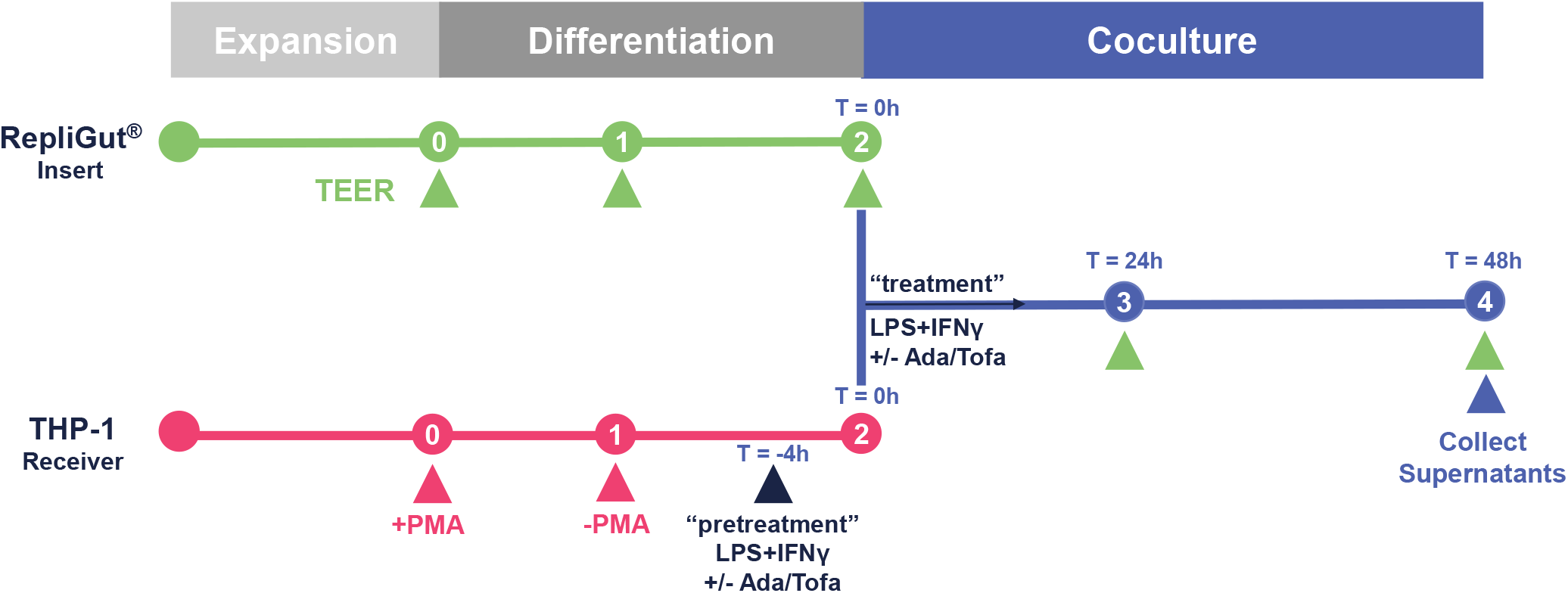
Timeline for RepliGut^®^ - Planar Transverse Colon primary human colonic epithelial cells and THP-1-dervied macrophages (THP-1m). <u>Transverse colon culture establishment (green timeline):</u> stem and progenitor cells are seeded on the cell culture inserts of a 96-well Transwell plate. Once confluent, TC cells were differentiated for 48 hours using RepliGut^®^ <u>Maturation Medium. Macrophage culture establishment (pink timeline):</u> On a separate receiver plate, THP-1 cells were seeded at 2.1 x 10^4^ cells/well (unless indicated otherwise) with medium containing PMA to induce differentiation. The following day, the medium was replaced without PMA for an additional 24 hours. For inflamed co-culture, THP-1m were pretreated with varying doses of LPS and IFN-γ 4 hours prior to initiation of co-culture. <u>Co-culture (blue timeline):</u> Differentiated TC monolayers in the apical compartment and THP-1m in the basal compartment were brought together and co-cultured for 48 hours. TEER was measured daily. For inflamed co-culture, treatment concentrations of LPS and IFN-γ were included in the apical and basal co-culture medium. Healthy groups were not treated with either LPS or IFN-γ.

**Figure 2.**
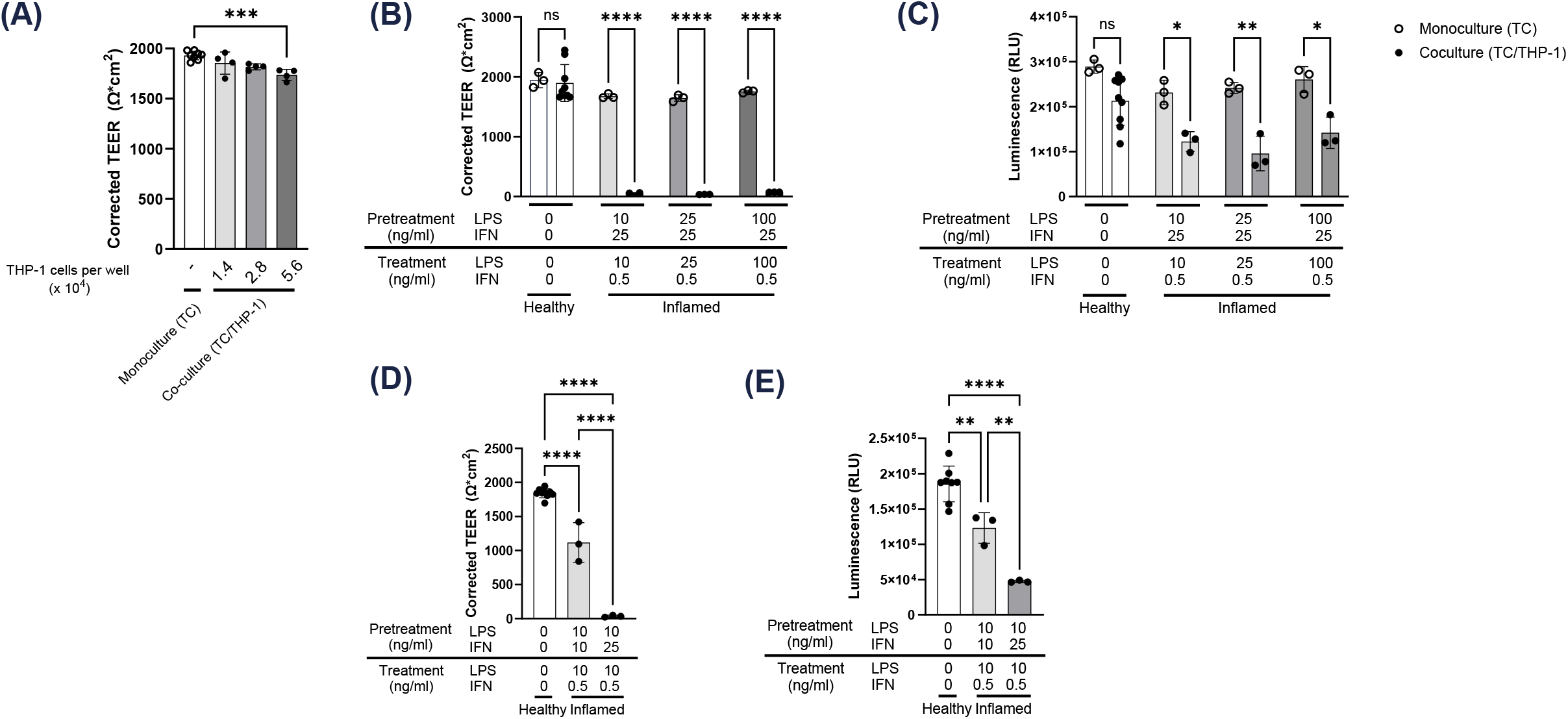
Optimization of THP-1 seeding density and stimulation regimen establish healthy and inflamed co-culture, assessed by barrier disruption and epithelial cell damage. **(A)** The effect of increasing seeding densities of THP-1 cells on epithelial barrier integrity was assessed using TEER in a healthy (unstimulated) state. Using 2.1 x 10^4^ THP-1 cells/well, cultures were treated with varying doses of LPS while holding IFN-γ concentrations constant: **(B)** TEER and **(C)** viability using Cell Titer Glo Assay were measured after 48 hours of co-culture. Using a constant dose of LPS, cells were treated with two doses of IFN-γ and TEER **(D)** and cell viability **(E)** were measured. Statistical analysis was performed using a Two-way ANOVA for B & C and One -way ANOVA for A, D, and E. * p < 0.05, ** p < 0.01, *** p < 0.001, **** p < 0.0001.

#### 2.1.3. Plating and Differentiation of RepliGut^®^ Planar Transverse Colon

Human intestinal tissues were obtained post-mortem from a certified organ procurement organization following ethical standards set by the Organ Procurement and Transplantation Network (OPTN; https://optn.transplant.hrsa.gov/). Donors were screened for HIV I/II, Hepatitis B (HBcAB, HBsAG), and Hepatitis C (HCV), and were confirmed to be healthy, without any known gut diseases. Intestinal crypts were isolated from the transverse colon, cultured under sub-confluent conditions, and cryopreserved as previously described (Pike et al., 2023; Wang et al., 2017). The donor cells used in this study were characterized as previously outlined (Pike et al., 2023).

For culture establishment, RepliGut^®^ - Planar Transverse Colon cell culture kits (Altis Biosystems, Cat#RGP-96W-TC) were used. Cryopreserved crypt-derived stem and progenitor cells were thawed in a 37 °C water bath and plated on proprietary matrix-coated Corning 0.4 µm 96-well Transwell^®^ plates in RepliGut^®^ Growth Medium (RGM). The apical and basal compartments contained 100 μl and 200 μl of medium, respectively. Once the cells reached confluence, the medium was switched to Altis RepliGut^®^ Maturation Medium (RMM) to induce polarization and differentiation for 48h before initiating co-culture with THP-1 cells. These steps correspond to the timeline steps shown in green in Fig. 1.

#### 2.1.4. Co-culture Establishment

These steps correspond to the timeline steps shown in blue in Fig. 1. Medium was aspirated from THP-1m and differentiated RepliGut^®^ - Planar Transverse Colon cultures, established as described above. Cells were washed once with 1X PBS. Cell culture inserts (containing differentiated transverse colon monolayers) were then transferred to the receiver plate containing THP-1m to initiate co-culture. The co-culture was maintained in 100 µl/200 µl apical/basal Co-culture Medium – THP-1 (CMT, Altis Biosystems, MED-CMT) for 48 hours. Control TC and THP-1m monocultures were also maintained in CMT from this timepoint onward.

#### 2.1.5. Co-culture Stimulation (Inflamed)

The THP-1m were pretreated with LPS from Escherichia coli K12 (LPS, Invivogen, Cat#tlrl-peklps) and IFN-γ protein (Peprotech, Cat#300-02) at the concentrations indicated in figure annotations for 4 hours in CMT, prior to initiation of co-culture. Medium was then removed, and cultures were combined as described in (d). During co-culture, CMT was supplemented with LPS at the concentrations indicated in figures and 0.5 ng/mL IFN-γ, along with either 3 µg/ml adalimumab (Ada., Selleck Chemicals, Cat#A2010), or 100 µM tofacitinib (Tofa., Selleck Chemicals, Cat#S2789). The co-culture was maintained for 48 hours. Apical and basal supernatants were collected at 48 hours following the initiation of co-culture and stored at -80°C for subsequent analysis.

### 2.2. Barrier Integrity: Transepithelial Electrical Resistance (TEER)

Barrier integrity was determined by measuring TEER on cultures daily using an EVOM™ Auto (World Precision Instruments EVA-MT-03-01). Raw TEER values for each well were corrected by subtracting pre-measured blank values and multiplying by well area in cm ^2^.

### 2.3. Cell Viability: Cell Titer Glo (CTG) Assay

Cell viability was assessed using CellTiter-Glo^®^2.0 (Promega, Cat#G9241) using the manufacturer’s instructions. Cell culture inserts were moved to a fresh receiver plate without THP1-m to assess cell viability separately from THP-1m cells. The reagent was mixed with an equal volume of CMT and added to the Transwell inserts or the original receiver plate. Following a 10 minute incubation at room temperature in the dark, the supernatant was collected from each well, and Luminescence was determined from two 40 μL samples from each well using a BioTek Synergy H1 plate reader. Individual data points on cell viability bar graphs represent the blank-subtracted average reading for duplicate samples from each well.

### 2.4. Cytokine Quantification using Luminex

Apical and basal cell culture supernatants were combined in a 1:1 ratio and analyzed for the following cytokines: TNFα, IL-1 beta, IL-6, IL-8(CXCL8), IL-12p70, IL-23; C-X-C Motif Chemokine Ligand 1 (CXCL11), and Granulocyte-Colony Stimulating Factor (G-CSF) (Invitrogen, Cat#PPX-08-MXWCYWA, Cat#EPX01A-10204-901(IL8)). The Luminex ProcartaPlex assays were performed following the manufacturer’s protocols, with blanks and standard curves included on each assay plate. Concentrations were determined of each analyte using Luminex™ xMAP™ Intelliflex.

### 2.5. Statistics

Statistical analyses were conducted using GraphPad Prism 9 (GraphPad Software, La Jolla, CA). All data are presented as the mean +/-standard deviation (SD), with data points representing individual cell culture replicates. One-way or Two-way Analysis of Variance (ANOVA) was performed, as indicated in individual figure legends, followed by Tukey’s post-hoc test for multiple comparisons. Statistical significance is indicated by asterisks in the figures: **p<0.01, *** p<0.001, ****p<0.0001.

## 3. Results

### 3.1 Development of Healthy Co-culture

The healthy co-culture model is designed to replicate a healthy intestine, where differentiated THP-1 macrophages (THP-1m) and primary transverse colon (TC) epithelial cells coexist without negatively affecting each other. Co-culture was initiated by placing apical inserts containing differentiated TC monolayers onto plates with THP-1m in the basal wells. Figure 1 depicts key aspects of the co-culture timeline for healthy and inflamed (stimulated with LPS and IFN-γ) states. Cultures were assessed for TC monolayer barrier function via TEER and cell viability using Cell-Titer Glo^®^ (CTG) after 48 hours of co-culture.

Developing a healthy baseline co-culture required optimization of the seeding density of THP-1 cells. High THP-1m densities may lead to stress and activation that causes release of inflammatory cytokines even in the absence of LPS and IFN-γ stimulation, which can compromise the epithelial monolayer barrier integrity (Kämpfer et al., 2017; Satsu et al., 2006). To optimize this, different seeding densities of THP-1 monocytes, ranging from 1.4 x 10^4^ to 5.6 x 10^4^ cells per well of a 96 well plate (surface area approximately 0.32 cm ^2^), were tested. At 48 hours of co-culture, we observed decreasing TEER correlated with increasing THP-1m density (Fig 2A). Notably, only the highest THP-1m density (5.6 x 10^4^ cells/well) resulted in a statistically significant difference as compared to healthy TC monocultures. A midpoint seeding density of 2.1 x 10^4^ cells/well was evaluated (data not shown) for its ability to generate sufficient cytokine release to drive barrier disruption via the conceptualized stimulation paradigm. This density not only supported baseline barrier integrity and viability in the absence of stimulation, as expected, but also was able to produce a desirable inflammatory response following stimulation. Following the initial positive outcome with 2.1 x 10^4^ cells/well, we proceeded with this density for subsequent experiments.

### 3.2 Development of Inflamed Co-culture

In the inflamed co-culture model, activated THP-1m are expected to secrete inflammatory cytokines that result in crosstalk with the TC epithelial monolayer, ultimately resulting in a measurable compromise to barrier integrity. To elicit an inflammatory response from THP-1m, we employed lipopolysaccharide (LPS), and interferon-gamma (IFN-γ) to induce M1 polarization and activation (Genin et al., 2015). To determine optimal concentrations for LPS + IFN-γ, we applied an initial LPS + IFN-γ pre-treatment cocktail (applied to THP-1m alone) in line with published values, followed by decreased IFN-γ in the LPS + IFN-γ cocktail applied during co-culture; concentrations of IFN-γ typically reported for THP-1m activation exceed the 3 ng/ml known to disrupt barrier integrity in RepliGut^®^ - Planar Transverse Colon cultures (data not shown). In an initial range-finding study, concentrations of lipopolysaccharide (LPS) up to 1000 ng/ml or interferon-gamma (IFN-γ) up to 1 ng/ml did not negatively impact the TC or THP-1m monocultures, as evidenced by sustained transepithelial electrical resistance (TEER) and cell viability (Fig. S1).

To determine the optimal concentrations of LPS and IFN-γ in combination for THP-1m pretreatment, we performed a dose-range study for LPS and IFN-γ. At t = 4 hours prior to co-culture, THP-1m were treated with 10, 25, or 100 ng/ml LPS in combination with 25 ng/mL IFN-γ. The same LPS concentrations were maintained during co-culture, while IFN-γ was reduced to 0.5 ng/ml. TC monocultures were treated identically to their corresponding co-culture groups. TEER of TC monolayers in vehicle-treated (“healthy”) co-cultures was not significantly different from TC in monoculture, as expected. TEER values remained high in TC monocultures exposed to all LPS + IFN-γ treatment concentrations tested, indicating preserved barrier integrity (Fig. 2B). In contrast, we observed near complete barrier disruption across all treated/inflamed co-culture conditions. Likewise, TC cell viability was not significantly different for TC in monoculture as compared to unstimulated co-culture (Fig. 2C). Viability remained high in TC monocultures exposed to all LPS + IFN-γ treatment concentrations, while TC cell viability was significantly decreased in the inflamed conditions. These results demonstrate achievement of measurable TC barrier disruption and cell viability reduction under the “Inflamed” treatment paradigm at all concentrations tested. Importantly, barrier disruption and reduced cell viability were achieved in the presence of THP-1m pretreated with LPS + IFN-γ, and not in response to the LPS + IFN-γ treatment conditions alone.

Given that the severity of TEER disruption and cell viability reduction was comparable for all LPS + IFN-γ combinations shown in Figure 2B & C, we further refined the IFN-γ dose to be used during the THP-1m pretreatment period (Fig. 2D, E). The LPS concentration was kept constant across pretreatment and treatment at 10 ng/ml, and we reduced IFN-γ pretreatment to 10 ng/ml. Co-treatment conditions remained constant. We observed a moderate response to 10 ng/ml IFN-γ pretreatment compared to 25 ng/ml with a partial drop in TEER (Fig. 2D) and viability (Fig. 2E).

### 3.3 Model Reproducibility

While the incorporation of primary cells promises to improve the relevance and accuracy of cell-based assays and microphysiological systems (MPS), they have classically faced challenges around reproducibility. Reproducibility of the inflamed state in our co-culture model is essential for its effective use. We believed that the most important source of variability to control in our system would be differences in TC cell lot. To characterize the reproducibility of the model across cell lots, we tested two lots from the same donor head-to-head, following the same dosing scheme used in Fig. 2D, E. Both cell lots exhibited a nearly identical 100% drop in TEER after 48 hours of co-culture under robust inflamed conditions, indicating a complete loss of barrier integrity. In contrast, healthy co-cultures maintained high TEER at 48 hours for both cell lots (Fig. 3A). The inflamed conditions exhibited reduced TC cell viability compared to the healthy vehicle control, indicating a statistically significant effect caused by the activated THP-1m (Fig 3B) in both TC cell lots. Furthermore, the results in Figure 3 alongside those in Figure 2 reflect good run-to-run consistency in the model. Data in Figure 2 was generated with “cell lot 1”; both runs show similar TEER and viability magnitudes and trends with stimulation, producing both a moderate and extreme damage to the epithelium. In combination, these results highlight the reproducibility of the inflamed co-culture model and confirm that THP-1m activation with the optimized LPS + IFN-γ dosing scheme reliably leads to significant epithelial barrier disruption.

**Figure 3.**
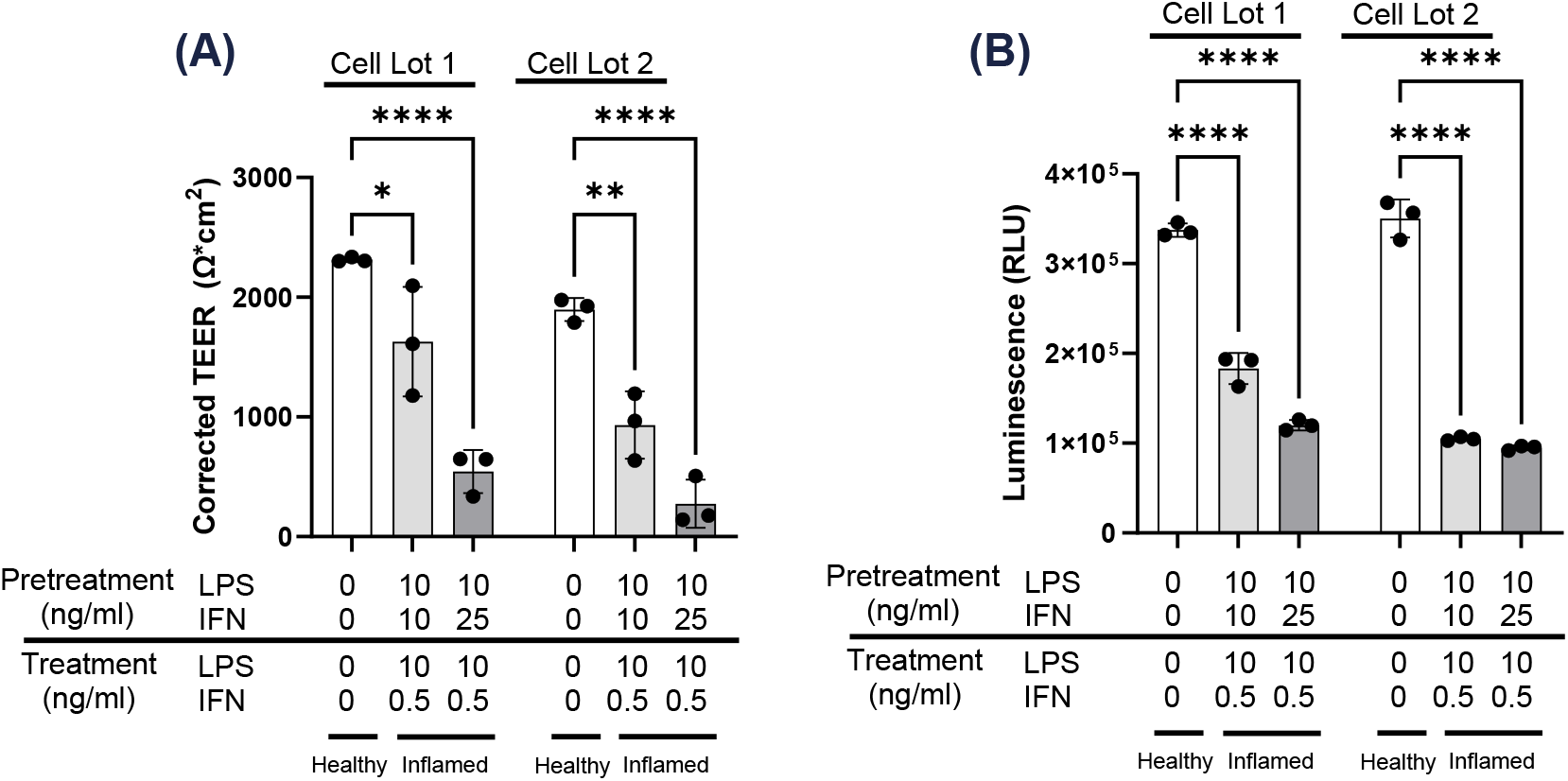
Barrier damage and cytotoxicity is reproduceable across epithelial cell lots in co-culture. To confirm reproducibility, two cell lots from the same donor were used to establish healthy and inflamed co-culture. THP-1m were pretreated with 10 ng/ml of LPS and either 10 ng/ml or 25 ng/ml of IFN-γ as performed in Figure 2 D & E. Treatment impact on **(A)** TEER and **(B)** viability was determined. Statistical analysis was performed using a Two-way ANOVA.* p < 0.05, ** p < 0.01, **** p < 0.0001.

### 3.4 Mitigating the Inflamed Condition Using Clinically Approved Drugs

To evaluate the model’s utility for screening anti-inflammatory drug and assessing their potential as clinical therapeutics, we employed two commercially available IBD drugs: adalimumab, a TNFα neutralizing antibody, and tofacitinib, a small molecule JAK inhibitor. We again performed this work in two different TC cell lots to further establish lot-to-lot reproducibility.

THP-1m were pretreated with LPS + IFN-γ in the presence of 3 µg/ml adalimumab (Ada) or 100 µM tofacitinib (Tofa). During co-culture, the medium was replaced with medium containing maintenance concentrations of LPS + IFN-γ and either Ada or Tofa for 48 hours. While LPS + IFN-γ stimulation led to a significant reduction in TEER and cell viability, the addition of Ada or Tofa prevented the damaging effects of LPS + IFN-γ treatment alone (Fig. 4); no significant difference was found in either cell lot or endpoint between Ada or Tofa treated inflamed groups and their corresponding healthy controls, as assessed by Tukey’s multiple comparisons following two-way ANOVA. Together, these data demonstrate that this model and readouts are effective at evaluating therapeutic interventions that target damaging cytokines produced by macrophages and epithelial cells in the gut.

**Figure 4.**
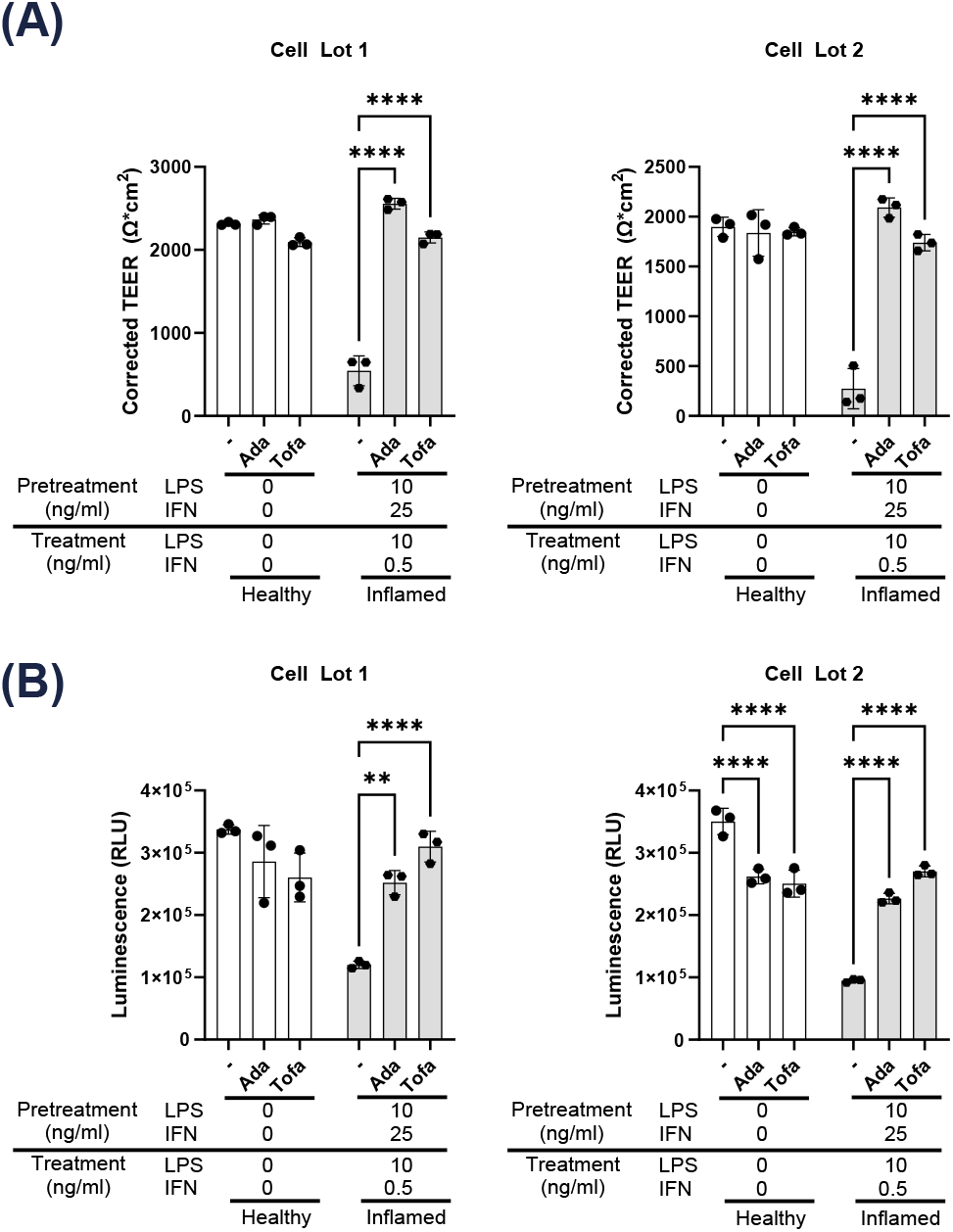
Epithelial cell damage can be mitigated by clinical therapeutics that target TNFα and JAK pathways. THP-1m were pretreated with LPS + IFN-γ in the presence or absence of 3 µg/ml of adalimumab (Ada) or 100 µM of tofacitinib (Tofa). These treatments were also carried into co-culture at the notated doses. TEER **(A)** and cell viability **(B)** were determined for two TC cell lots. Statistical analysis was performed using a Two-way ANOVA. ** p < 0.01, **** p < 0.0001.

### 3.5 Pro-Inflammatory Cytokine Profiling

Barrier disruption and cytotoxicity are reliable indicators of tissue damage. However, because the in vivo immune response involves complex signaling from additional immune cell types that are not present in the system, we measured a subset of proinflammatory cytokines released by macrophages and epithelial cells to capture relevant immune signaling. LPS + IFN-γ treatment drove release of pro-inflammatory cytokines (TNF-α, IL-1β, IL-6, IL-8, G-CSF, CXCL-11, and IL-23) in the inflamed condition compared to the healthy co-culture control (Fig. 5 A-G, Table S1). We also assessed IL-12 and did not observe a detectable difference (Table S1). Interestingly, adalimumab did not significantly reduce expression of most proinflammatory cytokines, except IL-8 in cell lot 1 (Fig 5D). Tofacitinib cotreatment significantly reduced all cytokine levels besides IL-23 (Fig. 5 and Table S1). Similar cytokine trends following stimulation with and without co-treatment were observed at the 10 ng/mL IFN-γ stimulation dose, though with reduced concentrations (Supplemental Table 1). These data confirm the release of a proinflammatory response that can be modulated by therapeutics, highlighting the model’s utility in functionally assessing cytokine activity.

**Figure 5.**
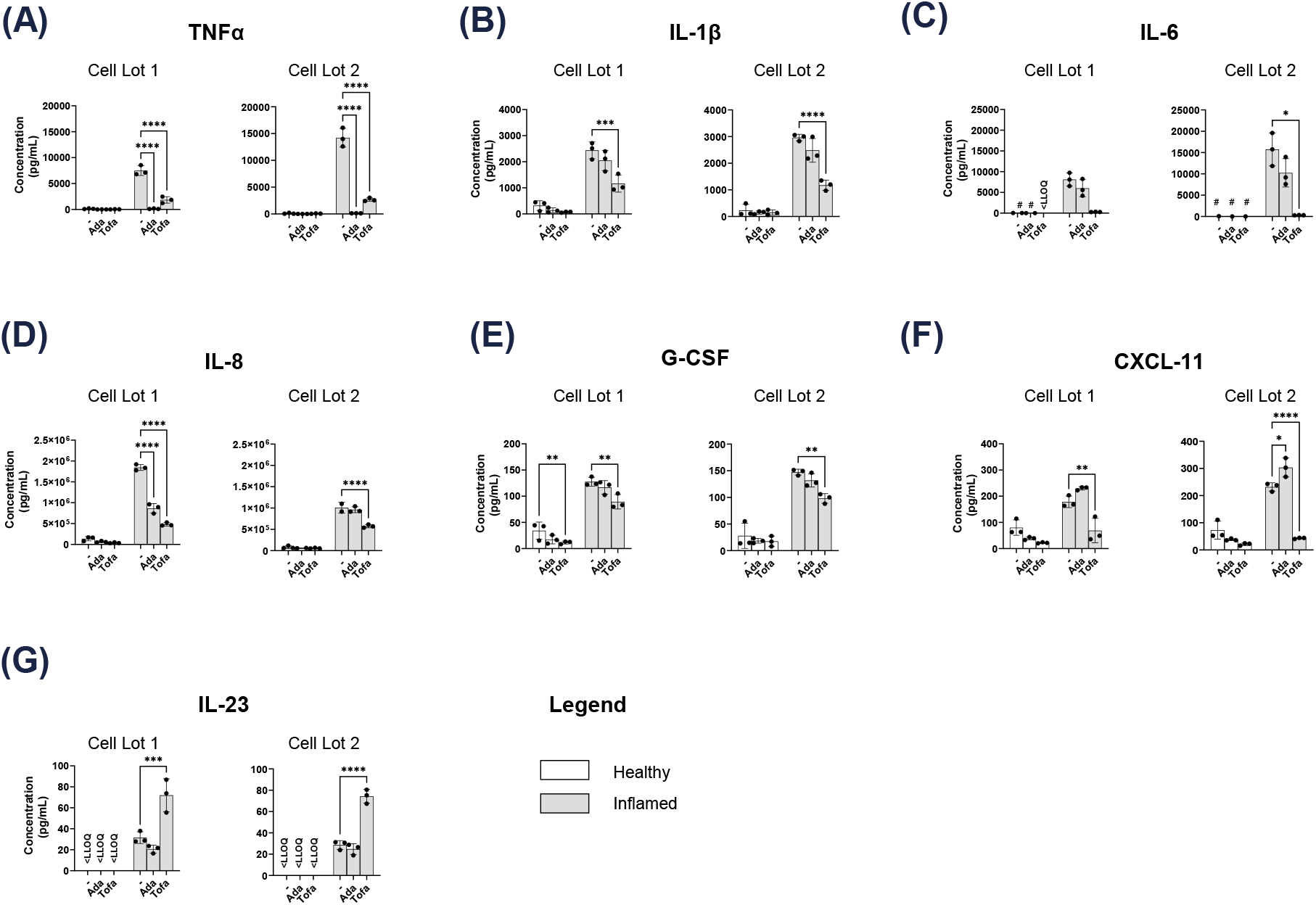
LPS and IFN-γ treatment induce a repeatable cytokine profile that is altered by clinical therapeutics. Cumulative proinflammatory cytokine release after 48 hours of co-culture was profiled for TC cell lots 1 and 2 in both healthy and inflamed conditions in the presence or absence of adalimumab (Ada) or tofacitinib (Tofa). Values below the lower limit of quantification (LLOQ) were excluded. # denotes groups with 1 or more data below LLOQ. Statistical analysis was performed using a Two-way ANOVA. * p < 0.05, ** p < 0.01, *** p < 0.001, **** p < 0.0001.

## 4. Discussion

Development of effective therapies for inflammatory bowel disease (IBD) is hampered by the limitations of preclinical models, which often fail to recapitulate the complex interplay between the human intestinal epithelium and the mucosal immune system. Animal models suffer from interspecies differences, while traditional *in vitro* models using immortalized cell lines like Caco-2 lack physiological relevance, particularly concerning epithelial differentiation, barrier complexity, and sensitivity to cytokines (Monticello et al., 2017; Olson et al., 2000; Sambuy et al., 2005). To address this gap, we successfully developed a high-throughput, human-relevant gut-immune co-culture model using primary human colonic epithelial cells (RepliGut^®^ -Planar Transverse Colon) and THP-1 derived macrophages. This co-culture system occupies a valuable niche: it offers improved physiological relevance over Caco-2-based co-cultures or primary monocultures, while maintaining well-characterized reproducibility and higher throughput than more complex fluidics-based MPS. The model effectively simulates both healthy intestinal homeostasis and key features of innate inflammation, including barrier disruption, reduced cell viability, and pro-inflammatory cytokine release, providing a robust platform for evaluating anti-inflammatory therapeutics.

A critical aspect for any screening platform is reproducibility. We demonstrated strong consistency between two independent lots of primary human colonic epithelial cells from the same donor. Both lots exhibited comparable baseline states, robust and consistent inflammatory responses (TEER, viability, cytokines) upon stimulation, and similar significant mitigation by adalimumab and tofacitinib (Fig 4, 5, Table S1). This lot-to-lot reproducibility significantly enhances confidence in the model’s reliability.

To establish an inflamed environment, we utilized LPS and IFN-γ to induce TLR4-mediated NF-κB activation and JAK/STAT signaling pathways, respectively. Together, these challenges have been previously optimized to promote macrophage M1 polarization and activation (Orecchioni et al., 2019; Pérez & Rius-Pérez, 2022). Our strategy of pretreating THP-1m with typical IFN-γ concentrations (25 ng/ml) reported in the literature, (Kämpfer et al., 2017) followed by a lower maintenance dose (0.5 ng/ml) during co-culture, successfully induced significant epithelial barrier disruption and viability reduction (Fig. 2, 3, 4) in the TC monolayers driven by activated macrophages rather than IFN-γ epithelial stimulation alone. As evidenced by the cytokine data (Fig. 5, Supplemental Table 1), a key strength of this model is the ability to induce an inflamed environment that includes release of cytokines commonly associated with IBD, giving researchers the ability to identify and study specific inflammatory signaling with endogenously produced cytokines. As expected for an M1-polarizing stimulus, we observed increased release of pro-inflammatory mediators compared to healthy controls. These included TNFα, IL-1β, IL-6, granulopoiesis promoter G-CSF, and the potent neutrophil chemoattractant IL-8 (CXCL8). Additionally, we observed increases in cytokines associated with T-cell infiltration and activation, such as T-cell chemoattractant I-TAC (CXCL11) and the Th1-promoting cytokine IL-23. The Th17-promoting cytokine IL-12 was below the lower limit of quantitation in all conditions (values reported in Supplemental Table 1). It should be considered that peak inflamed state concentrations of IL-23 were 32 pg/ml, while serum levels in humans range from 234 pg/ml (healthy) to 348 pg/ml (ulcerative colitis) (Mirsattari et al., 2012). Future work could explore alternative methods, stimulation doses for THP-1m activation or the addition of dendritic cells, to specifically drive additional secretion of Th1/Th17 axis cytokines. Together, cytokine characterization highlights the model’s utility in studying innate signaling pathways commonly associated with pathogenic challenge, as well as pathways associated with chronic inflammation, such as those seen in ulcerative colitis and Crohn’s disease.

The translational relevance of the model was demonstrated by its responsiveness to the clinically approved IBD therapies adalimumab and tofacitinib. As anticipated, the TNFα-neutralizing antibody adalimumab resulted in near-complete ablation of detectable TNFα (Fig. 5, Table S1), a decrease of downstream IL-8 with varying significance (Fig. 5), reduced barrier disruption, and increased cell viability compared to untreated inflamed conditions (Fig. 4). Neutralizing TNFα also led to decreasing trends in G-CSF, IL-1β, IL-6, and IL-12 (Supplemental Table 1), though these changes were not significant. Given that these cytokines are primarily produced by the THP-1m, it is unlikely adalimumab would reduce their secretion in the presence of continued IFN-y and LPS stimulation. TNFα produced by the THP-1m is expected to increase IL-8 release by both THP-1m and epithelial cells and increase epithelial cell cytotoxicity (Hideo et al., 2006), responses which were mitigated by adalimumab treatment. However, adalimumab treatment did not completely prevent IL-8 production in the system. Likely, the presence of LPS was enough to promote a level of IL-8 release via TLR4-induced Nf-kB activation. Future work could include TLR-4 antagonist in the system to further probe the IL-8 response. Nevertheless, the data clearly show that targeting TNFα dampens the overall inflammatory process and prevents cytokine-induced epithelial damage (Fig. 4 & 5).

Tofacitinib, a pan-JAK inhibitor targeting JAK1/3, demonstrated broad anti-inflammatory activity, significantly reducing levels of most measured cytokines including TNFα, IL-1β, IL-6, G-CSF, CXCL11, and IL-12 (Fig. 5, Table S1), while protecting barrier integrity and cell viability (Fig. 4). This effect is likely due to reduced activation of THP-1m in the system, which require IFN-y for activation in the current format (Taha et al., 2016). A counterintuitive finding emerged regarding IL-23: tofacitinib treatment consistently resulted in significantly **higher** levels of IL-23 compared to the inflamed vehicle control across both cell lots (Fig. 5, Table S1). This observation that tofacitinib increases IL-23 production above the inflamed control in our THP-1m co-culture is unexpected, given IL-23’s pro-inflammatory role. While paradoxical, it may stem from reducing anti-inflammatory signaling, such as IL-10 (Knoke et al., 2022). In vivo, IL-23 indirectly leads to epithelial disruption through the recruitment and activation of Th17 cells, which are not present in the current model. Despite the increase in IL-23 in this co-culture model system, tofacitinib exerted a strong net anti-inflammatory effect, suppressing key cytokines and robustly protecting against functional damage (TEER/viability).

This model incorporates primary colonic epithelial cells and THP-1-derived macrophages. While it does not include the full spectrum of immune cells - such as lymphocytes, dendritic cells, neutrophils, eosinophils, and innate lymphoid cells that are commonly associated with IBD (Velikova et al., 2024; Dwivedy and Aich, 2011) – it offers an easy-to-use, yet physiologically relevant platform for studying immune mechanisms specific to IBD. Future iterations could include additional intestinal regions of interest (e.g. the small intestine is commonly associated with Crohn’s disease) and/or an immune compartment that would enable exploration of adaptive responses in greater depth. This model system can be used to evaluate efficacy of anti-inflammatory drugs which act by distinct modes of action. Given that drugs in development for IBD and other autoimmune conditions have targeted nearly all of the cytokines we evaluated in this system or their receptors, the cytokine profiling underscores its broad utility for drug development and further enabling the discovery of novel therapeutic candidates and dissecting their impact on macrophage-epithelial crosstalk. It offers a significant advancement over traditional monocultures by incorporating key immune interactions, while retaining the throughput and reproducibility necessary for effective early-stage drug discovery, thereby helping to bridge the translational gap in IBD therapeutic development.

## 5. Conclusion

We have developed and characterized a reproducible, high-throughput human gut-immune co-culture model utilizing RepliGut^®^ Planar – Human Transverse Colon epithelia and THP-1 macrophages. This system successfully recapitulates key features of innate inflammation observed in IBD. An inflamed state, induced by activating macrophages with LPS and IFN-γ, led to the release of key inflammatory cytokines (including TNFα, IL-6, IL-1β) and subsequent epithelial barrier disruption. The model’s relevance for drug screening was confirmed by its responsiveness to clinically relevant anti-inflammatory drugs targeting different pathways: both adalimumab (anti-TNFα) and tofacitinib (JAK inhibitor) effectively mitigated inflammatory damage and maintained barrier integrity. Cytokine analysis revealed nuanced drug effects. This human-relevant model provides a valuable in vitro tool bridging the gap between overly simplistic cell cultures and complex animal studies, offering a promising platform to accelerate the discovery and development of novel IBD therapeutics.

## CRediT authorship contribution statement

*Swetha Peddibhotla:* Conceptualization, Methodology, Investigation, Validation, Data Curation, Visualization, Writing – Original Draft *Lauren A. Boone:* Investigation, Visualization, Writing – Review & Editing *Earnest Taylor:* Investigation, Visualization, Writing – Review & Editing *Bryan E. McQueen:* Conceptualization, Methodology, Writing – Review & Editing *Elizabeth M. Boazak:* Supervision, Project Administration, Conceptualization, Writing – Review & Editing.

## Acknowledgments

The authors thank Maureen Bunger and Jimmy Smedley for their valuable feedback and thoughtful review of the manuscript.

## Declaration of generative AI and AI-assisted technologies in the writing process

During the preparation fo this work, the authors used ChatGPT and Gemini 2.5 Pro iteratively with direct writing and editing by the authors to improve clarity and flow of the introduction and discussion. AI was also used to help generate reference suggestions. After using these services, the authors reviewed and edited the content as needed and take full responsibility for the content of the publication.

## Supplemental Figure Legends

**Supplemental Figure 1. Effect of LPS or IFN-γ alone on TC and THP-1m monocultures**. THP-1 cells were seeded at a density of 5.6 x 10^4^ cells/well. For Transverse Colon monocultures, cells were treated with or without LPS and IFN-γ for 48 hours and TEER **(A)** and viability **(B)** were assessed. For THP-1m monocultures, cells were pretreated with or without LPS at two concentrations (100 or 1000 ng/ml) or IFN-γ (25 ng/ml) for 4 hours. Subsequently, the THP-1m monocultures were treated again with LPS at the same concentrations or reduced IFN-γ concentrations. THP-1m viability was determined 48 hours after treatment **(B)**. Statistical analysis was performed using one-way ANOVA.

**Supplemental Table 1. Quantification of cytokines using Luminex**: Cumulative proinflammatory cytokine release after 48 hours co-culture was profiled for two TC cell lots in both healthy and inflamed conditions ± adalimumab (Ada) or tofacitinib (Tofa). Apical and basal supernatants were pooled 1:1 prior to analysis. The table shows cytokine values which are expressed as mean (pg/mL) + standard deviation and correspond to bar graphs in Figure 5. LLOQ = Lower Limit of Quantification.

