## Supplementary figures and images for "High-Throughput Human Gut Immune Co-Culture Model for Evaluating Inflammatory Bowel Disease Anti-Inflammatory Therapies"

### Supplemental_File_1

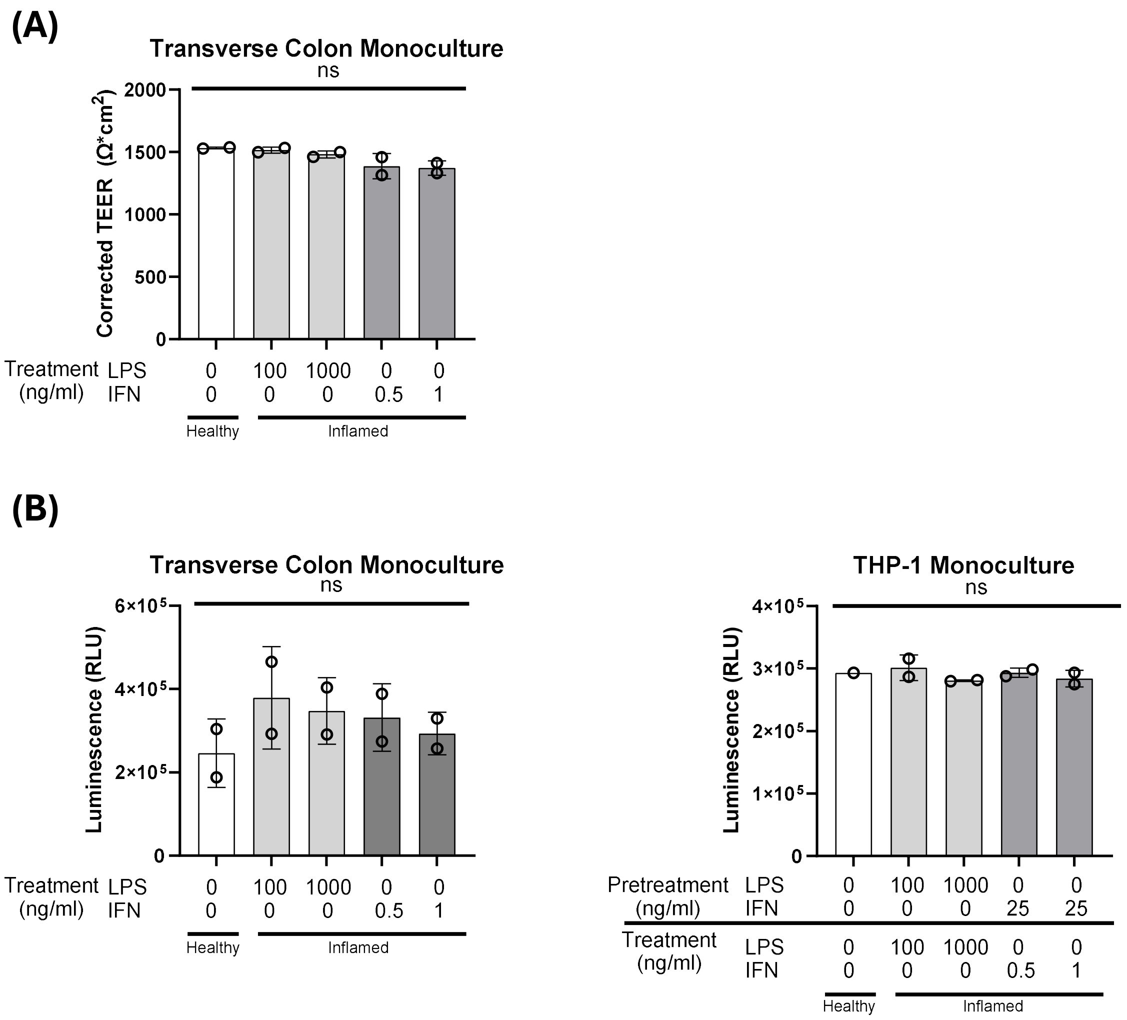
