## Supplemental_Table_1 for "High-Throughput Human Gut Immune Co-Culture Model for Evaluating Inflammatory Bowel Disease Anti-Inflammatory Therapies"

### Table S1.

LD0

Mean ± SD [pg/mL]

| | | Healthy | | | Inflamed (10+10) | | | Inflamed (10+25) | | |
|---|---|---|---|---|---|---|---|---|---|---|
| | | Vehicle | +Ada | +Tofa | Vehicle | +Ada | +Tofa | Vehicle | +Ada | +Tofa |
| Cell Lot 1 | TNFα | 137 ± 66 | 15 ± 7 | 30 ± 19 | 3,345 ± 1,174 | 150 ± 69 | 1,407 ± 445 | 7,492 ± 930 | 131 ± 60 | 1,856 ± 626 |
| Cell Lot 2 | | 99 ± 95 | 12 ± 3 | 64 ± 43 | 9,512 ± 651 | 137 ± 20 | 3,897 ± 979 | 14,228 ± 1,755 | 133 ± 5 | 2,667 ± 310 |
| Cell Lot 1 | G-CSF (CSF-3) | 34 ± 16 | 17 ± 8 | 12 ± 3 | 111 ± 5 | 103 ± 15 | 87 ± 13 | 128 ± 8 | 117 ± 13 | 89 ± 14 |
| Cell Lot 2 | | 28 ± 24 | 19 ± 5 | 18 ± 10 | 161 ± 8 | 156 ± 4 | 126 ± 12 | 147 ± 6 | 132 ± 13 | 98 ± 9 |
| Cell Lot 1 | I-TAC (CXCL11) | 80 ± 30 | 39 ± 6 | 22 ± 2 | 190 ± 10 | 208 ± 10 | 89 ± 70 | 178 ± 22 | 230 ± 6 | 69 ± 46 |
| Cell Lot 2 | | 73 ± 33 | 38 ± 4 | 21 ± 5 | 311 ± 28 | 376 ± 8 | 44 ± 6 | 232 ± 15 | 304 ± 35 | 43 ± 2 |
| Cell Lot 1 | IL-1β | 321 ± 184 | 141 ± 90 | 72±20 | 1,964 ± 306 | 1,583 ± 485 | 966 ± 268 | 2,437 ± 321 | 2,048 ± 390 | 1,164 ± 328 |
| Cell Lot 2 | | 231 ± 228 | 146 ± 57 | 147±96 | 3,866 ± 434 | 3,770 ± 368 | 2,209 ± 322 | 2,964 ± 120 | 2,484 ± 444 | 1,175 ± 193 |
| Cell Lot 1 | IL-6 | <LLOQ (26 ± 5) | <LLOQ (13 ± 5) | <LLOQ (0±0) | 6,618 ± 861 | 3767 ± 1,106 | 357 ± 145 | 9.759 ± 2,465 | 6,006 ± 2,060 | 294 ± 53 |
| Cell Lot 2 | | <LLOQ (35 ± 46) | <LLOQ (20 ± 0) | <LLOQ (8±6) | 17,675 ± 4,061 | 18,506 ± 5,629 | 556 ± 17 | 15,733 ± 3,877 | 10,250 ± 3,315 | 311 ± 33 |
| Cell Lot 1 | IL-8 (CXCL8) | 137,400 ± 50,619 | 63,725 ± 18,601 | 37,954±10,826 | 1,260,298 ± 123,698 | 983,254 ± 40,914 | 951,175 ± 48,614 | 1,844,349 ± 71,057 | 865,776 ± 113,191 | 481,337 ± 40,615 |
| Cell Lot 2 | | 81,359 ± 32,247 | 59,660 ± 4,309 | 57,514±9,381 | 750,685 ± 80,766 | 1,013,066 ± 84,573 | 790,522 ± 96,305 | 1,008,146 ± 127,053 | 961,194 ± 69,294 | 573,822 ± 37,380 |
| Cell Lot 1 | IL-12 | <LLOQ (0.3 ± 0.1) | <LLOQ (0.3 ± 0.1) | <LLOQ (0.3±0.1) | <LLOQ (3 ± 1) | <LLOQ (2 ± 0) | <LLOQ (2 ± 0) | <LLOQ (4 ± 1) | <LLOQ (3 ± 1) | <LLOQ (1 ± 0) |
| Cell Lot 2 | | <LLOQ (0.3 ± 0.1) | <LLOQ (0.3 ± 0) | <LLOQ (0.3±0.1) | <LLOQ (4 ± 1) | <LLOQ (4 ± 1) | <LLOQ (2 ± 0) | <LLOQ (5 ± 2) | <LLOQ (3 ± 1) | <LLOQ (1 ± 0) |
| Cell Lot 1 | IL-23 | <LLOQ (5 ± 1) | <LLOQ (5 ± 1) | <LLOQ (4±0) | 19 ± 3 | 20 ± 3 | 72 ± 23 | 32 ± 6 | 21 ± 4 | 72 ± 16 |
| Cell Lot 2 | | <LLOQ (4 ± 2) | <LLOQ (4 ± 1) | <LLOQ (4±1) | 23 ± 4 | 25 ± 2 | 129 ± 9 | 29 ± 4 | 25 ± 5 | 74 ± 7 |

#### Slide 1

**LD0** This is the same table as slide 11 but formatted differently.
Lauren Di Lella, 2025-05-09T13:50:40.182

**LD0 0** Make table editable
Lauren Di Lella, 2025-05-13T13:53:17.556
